# Long-read sequencing reveals the allelic diversity of the self-incompatibility gene across natural populations in *Petunia* (Solanaceae)

**DOI:** 10.1101/2024.06.18.599649

**Authors:** Taiga Maenosono, Kazuho Isono, Takanori Kuronuma, Miho Hatai, Kaori Chimura, Ken-ichi Kubo, Hisashi Kokubun, Julián Alejandro Greppi, Hitoshi Watanabe, Koichi Uehara, Takashi Tsuchimatsu

## Abstract

Self-incompatibility (SI) is a genetic mechanism to prevent self-fertilization and thereby promote outcrossing in hermaphroditic plant species through discrimination of self and non-self pollen by pistils. In many SI systems, recognition between pollen and pistils is controlled by a single multiallelic locus (called *S*-locus), in which numbers of alleles (called *S*-alleles) are segregating. Because of the extreme level of polymorphism of the *S*-locus, identifications of *S*-alleles have been a major issue in many SI studies for decades. Here we report an RNA-seq-based method to explore allelic diversity of the *S*-locus by employing the long-read sequencing technology of the Oxford Nanopore MinION, and applied it for the gametophytic SI system of *Petunia* (Solanaceae), in which the female determinant is a secreted ribonuclease called S-RNase that inhibits the elongation of self-pollen tubes by degrading RNA. We developed a method to identify *S*-alleles by the search of *S-RNase* sequences, using the previously reported sequences as queries, and found in total 62 types of *S-RNase* including 45 novel types. We validated this method through Sanger sequencing and crossing experiments, confirming the sequencing accuracy and SI phenotypes corresponding to genotypes. Then, using the obtained sequence data together with PCR-based genotyping in a larger sample set of 187 plants, we investigated the diversity, frequency, and the level of shared polymorphism of *S*-alleles across populations and species. The method as well as the dataset obtained in *Petunia* will be an important basis for further studying the evolution of S-RNase-based gametophytic SI systems in natural populations.

**Significance statement:** Flowering plants have evolved molecular mechanisms called self-incompatibility (SI) for discriminating self and non-self pollen at pistils to prevent self-fertilization, which is often deleterious due to inbreeding depression. The specificity of SI is usually determined by numbers of highly divergent alleles (called *S*-alleles) segregating at a single locus, and identifications of *S*-alleles have been a major issue in many SI systems. Here we report a new method to identify *S*-alleles by employing a long-read sequencing technology and applied it for the gametophytic SI system of *Petunia*, identifying 62 types of *S*-alleles including 45 novel types. The method as well as the dataset obtained in this study will be an important basis for the research of SI evolution.

## Introduction

Self-incompatibility (SI) is a genetic mechanism to prevent self-fertilization and thereby promote outcrossing in hermaphroditic plant species through discrimination of self and non-self pollen by pistils (De Nettancourt, 2001; Takayama and Isogai, 2005). SI systems are reported in more than 100 plant families and occur in about 40% of species (Igic et al., 2008). In many SI systems, recognition between pollen and pistils is controlled by a single multiallelic locus (called *S*-locus), in which numbers of alleles (called *S*-alleles) are segregating (Fujii et al., 2016; Takayama and Isogai, 2005). On the *S*-locus, a female specificity gene and a male specificity gene are tightly linked, and pollen is rejected when the same specificity of the *S*-allele is expressed by both pollen and pistils. There are two major distinct forms of SI mechanisms, the sporophytic SI (SSI) and the gametophytic SI (GSI), which differ by the genetic determination of the pollen specificity phenotype (Takayama and Isogai, 2005).

In most of the SI systems reported, the *S*-locus shows an extreme level of polymorphism (Schierup and Vekemans, 2008). Numbers of *S*-alleles are segregating in a population and *S*-allele sequences are highly divergent from each other (Castric and Vekemans, 2004). Such high allelic and sequence variation at the *S*-locus reflects negative frequency-dependent selection, where pollen of rare *S*-alleles is rejected by pistils at lower rates than those of common *S*-alleles (Castric and Vekemans, 2004; Schierup and Vekemans, 2008; Vekemans and Slatkin, 1994; Wright, 1939). Identifications of *S*-alleles have been a major issue in many studies of SI systems for decades, because they are a fundamental basis for studying the genetic and evolutionary features of SI, including the number of segregating *S*-alleles in populations, associations between *S*-alleles and SI phenotypes, trans-specific sharing of polymorphism, detecting natural selection on the *S*-locus genes, and *S*-allele distribution across populations (Castric and Vekemans, 2004; Durand et al., 2020).

However, the identification and genotyping of *S*-alleles have been challenging because of their extreme level of polymorphism. Since the male and female specificity genes are identified in multiple SI systems (Fujii et al., 2016; Takayama and Isogai, 2005), molecular cloning and Sanger sequencing of S-alleles have been major approaches for identification. These methods require polymerase chain reaction (PCR) and general primers designed in conserved regions intended to amplify specificity-determining genes irrespective of *S*-alleles (Charlesworth et al., 2000; Mable et al., 2003). One of the major obstacles to these approaches is that a set of general primers is rarely perfect, often failing to amplify some of the *S*-alleles, particularly highly divergent ones. It has also been suggested that PCR would amplify closely related sequences that are not linked to the *S*-locus (Mable et al., 2003).

Recently, next-generation sequencing technologies have been applied for the identification and genotyping of *S*-alleles. (Jørgensen et al., 2012; Mable et al., 2018, 2017) adopted a barcoded amplicon-based method and successfully identified *S-*alleles in multiple *Arabidopsis* species, although these methods were sensitive to biases of PCR amplification. (Tsuchimatsu et al., 2017) exploited a short-read-based whole-genome resequencing data of 1,083 natural accessions of self-compatible *A. thaliana* and revealed allelic variation of the *S*-locus. This method was based on a mapping to reference sequences available for all three *S*-alleles segregating in *A. thaliana*, thus would not directly be applicable to the identification of many novel *S*-alleles in self-incompatible species. More recently, (Genete et al., 2020) developed an integrated pipeline to predict *S*-alleles from short-read data by combining mapping-based and de novo assembly approaches, successfully identifying previously reported *S*-alleles as well as novel ones in *Arabidopsis halleri*. These methods using next-generation sequencing data were so far mostly designed and applied for SSI systems such as those of *Arabidopsis*, rarely for GSI systems. While (De Franceschi et al., 2018) was an example proposing a resequence-based approach for the GSI system of Rosaceae, their method was intended to gain full-length coding sequences rather than obtaining novel *S*-allele sequences from large-scale samples.

Here we report an RNA-seq-based method to explore allelic diversity of the *S*-locus by employing the long-read sequencing technology of the Oxford Nanopore MinION, and applied it for the GSI system of *Petunia* (Solanaceae). In this system, the female determinant is a secreted ribonuclease called S-RNase, inhibiting the elongation of self-pollen tubes by degrading RNA (Takayama and Isogai, 2005). The male determinant is a set of F-box proteins, called S-locus F-box (SLF), functioning as a component of the SCF (Skp1–Cullin1–F-box)-type E3 ubiquitin ligase that generally mediates ubiquitination of target proteins for degradation by the 26S proteasome (Takayama and Isogai, 2005). This system has been considered a collaborative non-self recognition system, in which the product of each SLF interacts with a subset of non-self S-RNases, and the products of multiple SLF types are required for the entire suite of non-self S-RNases to be recognized (Fujii et al., 2016; Kubo et al., 2010, 2015). In this study, we first developed a pipeline to identify *S*-alleles based on the sequences of *S-RNase*, using the previously reported sequences as queries (Fig. 1). MinION-based RNA-seq enabled us to directly obtain full-length transcripts of *S-RNase*, avoiding the possibility of misassembly that could occur with the de novo assembly of short-read sequences. Next, we validated this method through Sanger sequencing and crossing experiments. Then, using the obtained sequence data together with PCR-based genotyping in a larger sample set, we investigated the diversity, frequency, and distributions of *S*-alleles across populations and species. The method as well as the dataset obtained in *Petunia* will be an important basis for further studying the evolution of non-self-recognition systems in natural populations.

**Fig. 1.**
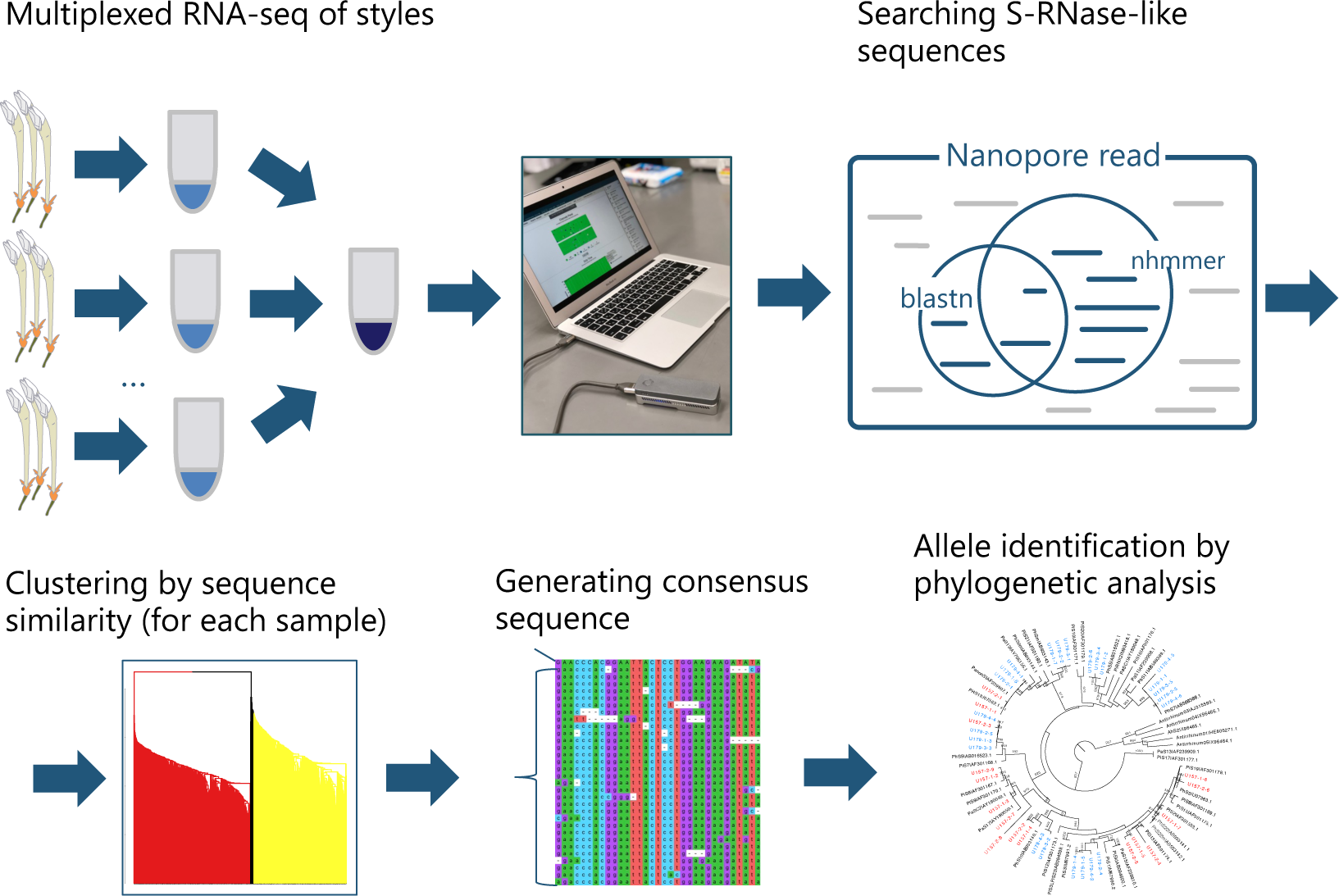
Overview of the developed pipeline to identify *S-RNase* sequences. We performed individually barcoded multiplexed RNA-seq of styles using the Nanopore MinION sequencer, and searched for S*-RNase*-like sequences using previously reported *S-RNase* sequences as queries. Then, two heterozygous *S-RNase* sequences were separated by clustering analysis based on sequence similarity, and consensus sequences were generated for each *S*-allele. Finally, we generated a phylogenetic tree and assigned *S*-alleles based on the monophyly.

## Results

### Identification of *S-RNase* by Nanopore sequencing

Our developed pipeline is summarized in Fig. 1. In short, we performed individually barcoded multiplexed RNA-seq of styles, and searched for *S-RNase*-like sequences using previously reported *S-RNase* sequences as queries. Then, two heterozygous *S-RNase* sequences were separated by clustering analysis, and consensus sequences were generated for each *S*-allele. Finally, we generated a phylogenetic tree and assigned *S*-alleles based on the monophyly.

We first performed RNA-seq of styles using the Oxford Nanopore MinION sequencer for 86 individuals in total (Table S1). After trimming adaptor sequences, mean number of reads and total base pairs per sample were 219,078 and 85,739,454 bp, respectively (Table S1). Using BLAST (v.2.2.31) (Altschul et al., 1990) and HMMER (v.3.3.2) (Finn et al., 2011), we searched for *S-RNase*-like sequences using publicly available 37 S-RNase sequences as queries, and found on average 654 *S-RNase*-like reads per sample, ranging from 13 to 5994 (Table S2). In each individual, we separated those *S-RNase*-like sequences by clustering analysis based on sequence similarity, and obtained a consensus sequence for each cluster. After excluding two reported *S-RNase*-like sequences unlinked to self-incompatibility (*Panon-S* and *PiRNX2*; (Lee et al., 1992)), we in total obtained 172 sequences and generated a phylogenetic tree together with previously reported *S-RNase* sequences of *Petunia*, and those of *Antirrhinum* as outgroups (Fig. 2A). We then assigned *S*-alleles based on the monophyly of this tree, identifying 62 alleles including 45 novel *S*-alleles. It is important to note that, except for the alleles unlinked to self-incompatibility (*Panon-S* and *PiRNX2*), we found two *S*-alleles copies from all the individuals surveyed, suggesting the validity of our approach (Table S2).

**Fig. 2.**
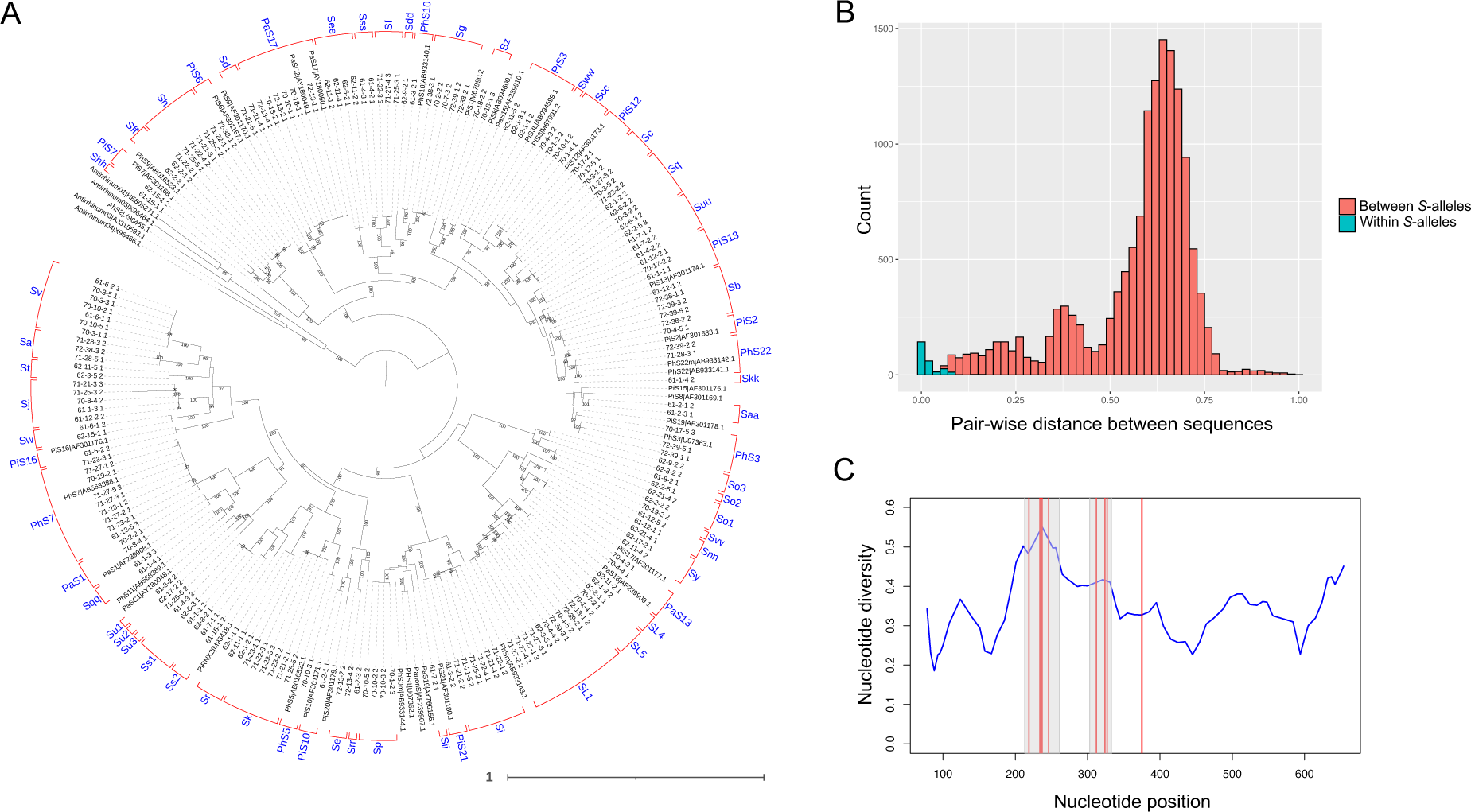
Identification of *S-RNase* sequences in *Petunia*. (A) A maximum likelihood phylogenetic tree of the *Petunia S-RNase* sequences with *Antirrinum* ones as outgroups. The percentage of 1,000 bootstrap replicates are shown on the branches when ≥ 90%. Assigned names of *S*-alleles are also shown in blue. (B) Pair-wise genetic distance between *S-RNase* sequences. Distances between different and the same *S*-allele pairs are indicated by red and blue, respectively. (C) Nucleotide diversity along the *S-RNase* genes (blue line) and positively selected sites (red vertical lines). Gray shaded areas indicate hypervariable regions I and II.

Fig. 2B shows the distribution of pair-wise genetic distances between all the *S-RNase* sequences in the tree with the information of our *S*-allele assignments. While our *S*-allele assignments are not strictly based on the genetic distance between sequences but on the monophyly in the tree, this histogram shows that the two *S*-alleles were largely assigned as the same ones if their genetic distance is in the left-most peak, suggesting that our *S*-allele assignments have consistency with the genetic distance between *S*-alleles.

To evaluate the sequencing accuracy of Nanopore-based genotyping, we performed Sanger sequencing for in total 69 *S*-alleles and compared them with Nanopore-based sequences (Table S3). We found that the sequencing accuracy was 99.1% (median value), suggesting that the Nanopore-based sequencing is mostly accurate and reliable.

Using these sequences obtained by the Sanger sequencing method, we evaluated the genetic diversity along the *S-RNase* gene by sliding window analysis (Fig. 2C). While the nucleotide diversity of the gene was overall high (meanπ= 0.341), it was particularly elevated in the hypervariable regions I and II as previously reported (Ioerger et al., 1991). Since high genetic diversity is indicative of positive selection, we formally detected sites under positive selection in *S-RNase* using the PAML4 software with the Bayes empirical Bayes method (Yang, 2007). The likelihood of the model M2a (positive selection) was significantly higher than the model M1a (nearly neutral) (likelihood ratio test, *P* = 1.78 x 10^−24^), indicating that several nucleotide positions are under positive selection. Nine positive selection sites with a posterior probability > 0.95 were identified, and eight of them were located in the hypervariable regions I or II (Fig. 2C).

### Frequency and distribution of *S*-alleles

To investigate the frequency and distribution of *S*-alleles in a larger sample set, we performed PCR-based genotyping for individuals that were not used for Nanopore sequencing. We first tested the validity of the PCR-based genotyping by Sanger sequencing of several individuals that were assigned as the same *S*-alleles (*S*v and *S*g alleles; Fig. S1). Both trees show that individuals assigned to harbor the same *S*-alleles by PCR-based genotyping had almost identical sequences, demonstrating the validity of PCR-based genotyping.

We performed genotyping of in total 187 individuals from three populations of *P. axillaris* subsp. *axillaris* and two populations of *P. inflata* (Tables S4 and S5). Note that our sample set of 187 individuals was comprised of half-sibs of 40 families. Among in total 374 copies of 187 individuals, we could not identify 20 copies (5.3%), suggesting the presence of unknown alleles.

We then investigated the frequency and distribution of *S*-alleles across populations and species (Fig. 3). The frequency of *S*-alleles varied greatly, ranging from 0.28% (singleton) to 4.8%, although isoplethy is theoretically expected in GSI systems (Wright, 1939). The number of *S*-alleles per population ranged from 9 to 27. Among the 62 *S*-alleles identified, 21 *S*-alleles (33.9%) were found in more than one population and 14 *S*-alleles (22.6%) were found in two species, suggesting the pervasive shared polymorphism between populations and species.

**Fig. 3.**
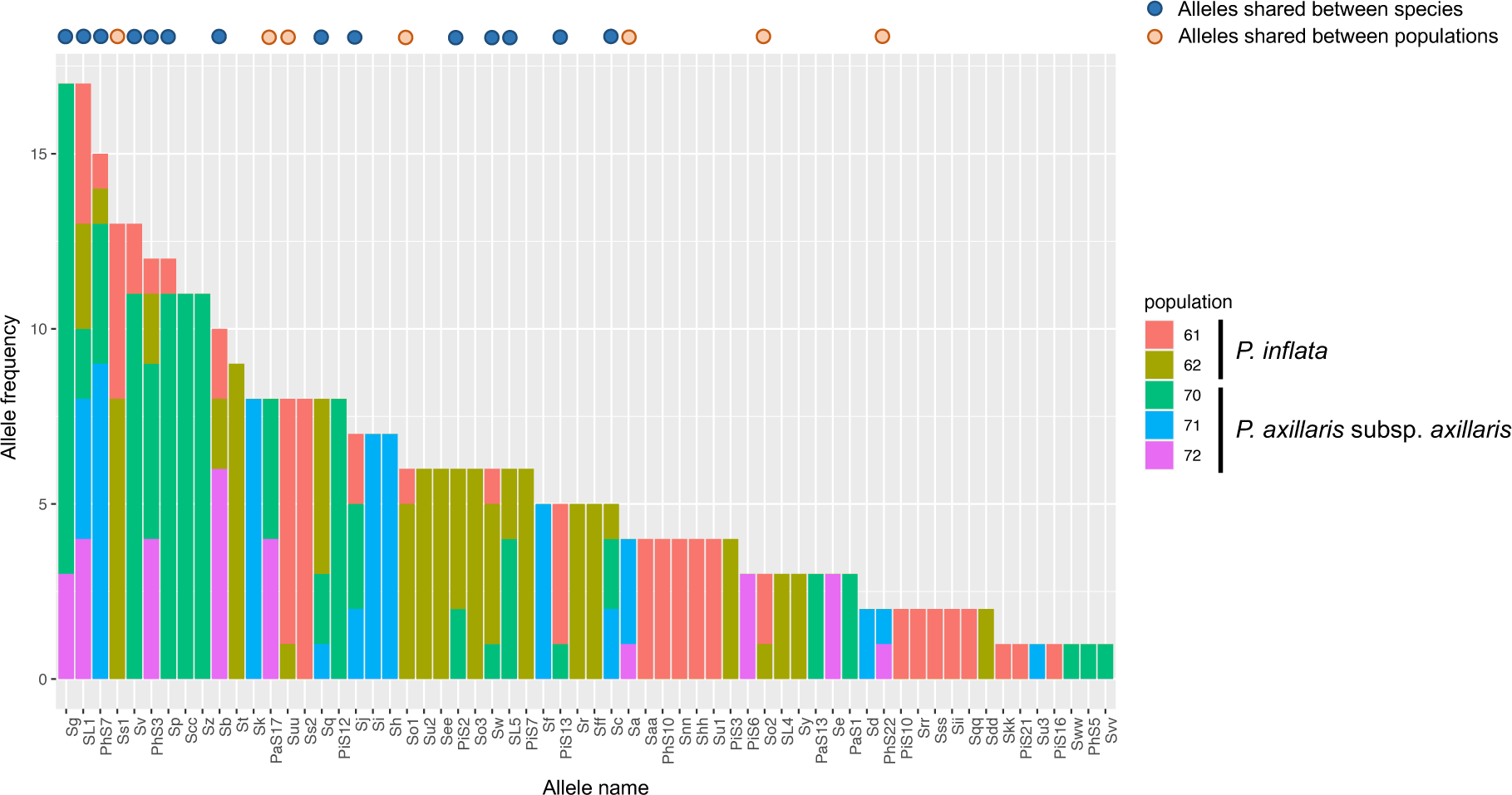
Frequency and distribution of *S*-alleles. Frequencies of each population are shown for each *S*-allele. The data is based on the genotyping of in total 187 individuals from three populations of *Petunia axillaris* subsp. *axillaris* and two populations of *Petunia inflata*. *S*-alleles shared between species or between populations are indicated on the top of the histogram.

The total number of *S*-alleles was estimated for each population using two different methods: the *E*_2_ estimator (O’Donnell and Lawrence, 1984), and a curve fitting with a two-parameter Michaelis–Menten model (Busch et al., 2010). The former ranged from 11 to 31, and the latter from 12 to 41, both of which were higher than the actual numbers of *S*-alleles observed (Table 1). This result suggests that we could not exhaustively identify all the *S*-alleles segregating in each population, consistent with the presence of unknown *S*-alleles in our PCR-based search of 187 individuals (Table S4).

**Table 1.** Observed and estimated numbers of *S*-alleles in five populations.

| Species | Population code | Number of copies investigated | Observed number of *S*-alleles | O'Donnell and Lawrence (1984) | Busch et al. (2010) |
| --- | --- | --- | --- | --- | --- |
| *Petunia axillaris*<br>subsp. *axillaris* | 70 | 105 | 22 | 25.56 | 27.74 |
| *Petunia axillaris*<br>subsp. *axillaris* | 71 | 52 | 13 | 14.68 | 16.74 |
| *Petunia axillaris*<br>subsp. *axillaris* | 72 | 29 | 9 | 11.82 | 12.15 |
| *Petunia inflata* | 61 | 74 | 27 | 31.60 | 41.39 |
| *Petunia inflata* | 62 | 94 | 24 | 27.97 | 31.09 |

To understand how *S*-alleles are shared across genera in Solanaceae, we constructed a phylogenetic tree by using *S-RNase* sequences of *Petunia* and other genera, *Physalis, Iochroma, Eriolarynx, Lycium, Solanum, Brugmansia, Vassobia, Dunalia, Nicotiana, and Witheringia*, in total 646 sequences, including *Antirrhinum* as outgroups. We found that the tree was not strictly clustered by genera, but many *S*-alleles were shared between genera (Fig. 4). We also found some clades with only specific genera, suggesting recent genus-specific expansions.

**Fig. 4.**
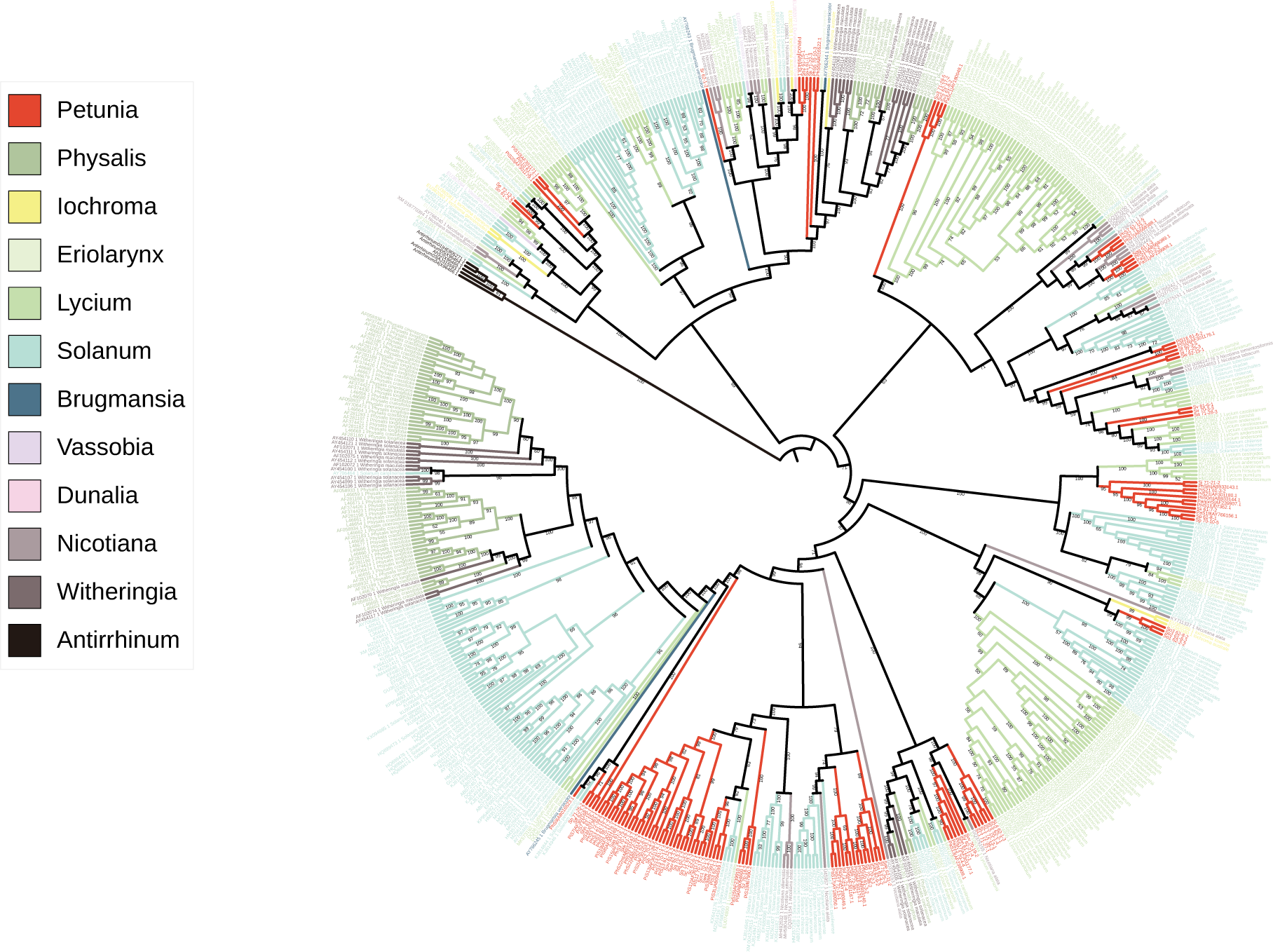
A maximum likelihood phylogenetic tree of *S-RNase* using 646 sequences from *Petunia* and other genera of Solanaceae, *Physalis, Iochroma, Eriolarynx, Lycium, Solanum, Brugmansia, Vassobia, Dunalia, Nicotiana,* and *Witheringia*, in total 646 sequences, including *Antirrhinum* as outgroups. The values on the branches indicate the percentage of 1,000 bootstrap replicates.

### Genotypes correspond to phenotypes

Our *S*-allele assignments based on sequence similarity may not necessarily correspond to SI specificities, and thus it is important to show the link between genotypes and the self-incompatibility phenotypes. We therefore performed crossing experiments using *P. axillaris* subsp. *axillaris* individuals and observed self-incompatibility phenotypes using in total nine *S*-alleles (Fig. 5; Table S6). We first performed crosses between seven *S*-alleles (*S*7, *S*22, *S*17, *S*3, *S*g, *S*f and *S*j) and overall observed the incompatibility phenotype in crosses between individuals that have the same *S*-alleles but observed the compatible phenotype in crosses between individuals that have different *S*-alleles (Fig. 5A; Table S6). We also performed crosses between closely related *S*-alleles (*S*6, *S*17, and *S*d) and consistently observed compatible reactions in all outcrosses (Fig. 5B), suggesting that the SI specificities are already differentiated in these closely related *S*-alleles. It is also suggested that our assignment of *S*-alleles based on sequence similarity would largely reflect the SI specificities, demonstrating the validity of our method.

**Fig. 5.**
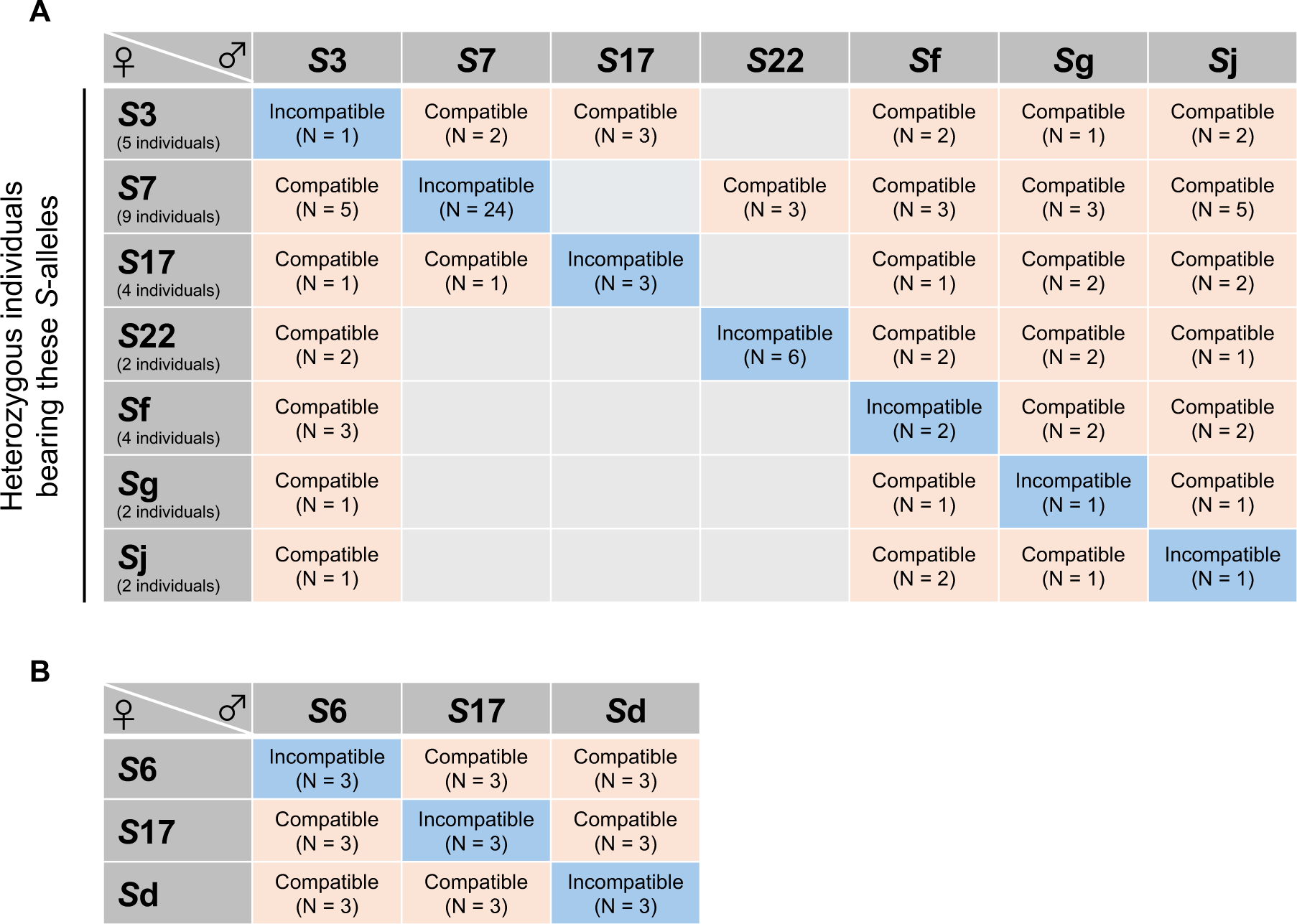
Crossing experiment to examine whether self-incompatibility phenotypes are associated with S-RNase genotypes. Individuals of *Petunia axillaris* subsp. *axillaris* were used for experiments. We observed pollen tubes by aniline staining and less than 20 pollen tubes reaching the bottom tip of styles were considered as a criterion of incompatible crosses. (A) Seven homozygous lines generated by forced selfing were used as pollen donors (*S*7, *S*22, *S*17, *S*3, *S*g, *S*f, and *S*j), and heterozygous individuals that have at least one of these *S*-alleles were used as pistil donors (see Table S6 for details). (B) Homozygous lines of *S*6, *S*17, and *S*d alleles were used as pollen and pistil donors. Note that these three *S*-alleles (*S*6, *S*17, and *S*d) had closely related *S-RNase* sequences.

## Discussion

### A MinION-based method to detect *S-RNase* alleles

In this study, we developed an RNA-seq-based method to identify and genotype *S-RNase* genes exploiting the long-read sequencer, MinION. We identified 62 alleles including 45 novel *S*-alleles through the sequencing of 86 individuals (172 copies). Based on the obtained sequence information, we further genotyped 187 individuals by PCR to investigate the *S-*allele distributions and frequencies in a larger sample set.

The validity of our approach was supported by multiple lines of evidence. First, in the individuals used for MinION-based RNA-seq, we identified two *S*-alleles per diploid individual, when previously reported non-specific allele copies (*Panon-S* and *PiRNX2*) were excluded. The PCR genotyping based on these MinION sequences also identified two *S*-alleles per individual (Table S4). These results suggest that our method could overall identify true *S*-allele sequences properly rather than merely list up *S-RNase*-like sequences. Second, crossing experiments based on the genotypes of *S-RNase* confirmed that the SI phenotypes were largely correlated with genotypes (Fig. 5; Table S6). The result supports the validity of our method to assign *S*-alleles based on the *S-RNase* sequence similarity.

Several methods have been used to genotype the *S-RNase* gene in GSI systems, such as two-dimensional gel electrophoresis (Sassa et al., 1994), restriction enzyme-based genotyping (cleaved amplified polymorphic sequences and restriction fragment length polymorphism) (Kim et al., 2009; Takasaki et al., 2004), and PCR with allele-specific or general primers (Broothaerts, 2003; Dzidzienyo et al., 2016; Janssens et al., 1995; Kubo et al., 2010, 2015; Matsumoto et al., 2006, 1999; Richman et al., 1995; Robbins et al., 2000; Sheick et al., 2020; Verdoodt et al., 1998), but false positives, false negatives, and the throughput of the experiments have been major obstacles. We demonstrate that our MinION-based method can identify novel *S*-alleles and detect known *S*-alleles in *Petunia*, which would be applicable to other S-RNase-based GSI systems.

Despite the validity of our method to detect *S-RNase* alleles, here we note a few caveats. First, although we confirmed MinION-generated sequences through Sanger sequencing, sequences were not perfectly matched in multiple *S*-alleles. Therefore, while our method has enough power to discover novel *S*-alleles and genotype them, it would be desirable to confirm by Sanger sequencing for the complete determination of the nucleotide sequences. Second, this method relies on the relatively high expression levels of the *S-RNase* gene in pistils, which provide hundreds of reads and help reduce sequence errors. Therefore, this method may not be applicable when the expression level of *S-RNase* is low (e.g. self-compatible strains). Third, our method benefited from the relatively abundant available *S-RNase* sequences of *Petunia* as queries, thus it may not work as efficiently as this study in taxa with little *a priori* information of *S-RNase*.

### Allele number, distribution, and frequency

Based on the genotyping data of 187 individuals, we investigated the frequencies of *S*-alleles and their distributions across five populations from two species, *P. axillaris* and *P. inflata*. We estimated the total number of *S*-alleles for each population using two methods (Busch et al., 2010; O’Donnell and Lawrence, 1984) The former ranged from 11 to 29, and the latter from 12 to 41, both of which were higher than the actual numbers of *S*-alleles observed (Table 1). We note that these should still be underestimates because our sample set consists of half-sib families. They are more likely to share the same *S*-alleles between individuals than random sampling, which the applied models assume.

In this study, among the 62 *S*-alleles identified, 21 *S*-alleles (33.9%) were found from more than one population. (Richman et al., 1995) investigated *S*-alleles in two wild populations of *Solanum carolinense*, and found 11 and 12 *S*-alleles for each, 10 of them were shared between populations. A study using *Prunus lannesiana* var. *speciosa* investigated *S*-alleles in seven populations, finding that *S*-alleles are shared 41-83% (Kato et al., 2007). These differences in the fraction of shared *S*-alleles may be due to the sample size, the genetic divergence between populations, and the classification of *S*-alleles. If our assignment of *S*-alleles based on the sequence similarity does not correspond to the SI specificity in some cases, the fraction of shares *S*-alleles between populations may be over- or under-estimated. For example, a few high-frequency *S*-alleles found in all five populations (e.g., *S*L1) showed population-specific clades (Fig. 2A), thus it might be possible that *S*-allele specificities are already differentiated between populations. In such a case, the fraction of shares *S*-alleles is overestimated.

### Toward an understanding of *S*-allele evolution

There have been extensive studies of the S-RNase-based SI system in *Petunia*, as the collaborative non-self recognition system was first discovered in this genus. Despite the wealth of knowledge of the functional aspect of the S-RNase-based SI system, the comprehensive information on *S*-locus diversity in wild populations of *Petunia* remained largely unexplored. Indeed, while about 40 *S-RNase* partial or full-length sequences have been reported so far, many of them were obtained from cultivated species *Petunia* x *hybrida* or a relatively limited number of strains in wild populations (Ai et al., 1992, 1990; Clark et al., 1990; Clark and Sims, 1994; Coleman and Kao, 1992; Entani et al., 1999; Kubo et al., 2010, 2015; Sassa and Hirano, 2006; Tsukamoto et al., 2005, 2003; Wang et al., 2001, 2003). Among the S-RNase-based GSI systems, the survey of *S*-allele diversity in wild populations is still limited to a handful number of species, including *Solanum carolinense* (Richman et al., 1995), *Solanum chilense* (Igic et al., 2007), *Sorbus aucuparia* (Raspé and Kohn, 2007), *Prunus lannesiana* var. *speciosa* (Kato et al., 2007), and *Malus sieversii* (Ma et al., 2017). One of the major questions in this field would be to demonstrate how a new specificity of SI evolves in the non-self recognition system. Although there are several theoretical investigations on the origin of new *S*-alleles (Bod’ová et al., 2018; Fujii et al., 2016; Harkness and Brandvain, 2021; Kubo et al., 2015), the information in wild populations is essential, including the number of *S*-alleles, variation in *SLF* repertories between *S-*alleles. Here we laid the foundation for the studies of the evolution of SI specificities in the non-self recognition system by surveying *S*-alleles in the wild *Petunia* populations. In particular, the recently diverged S-RNase sequence pairs would serve as a good system to study the origin of new SI specificities, as recently demonstrated in the SSI system (Chantreau et al., 2019).

## Materials & Methods

### Plant materials

We used in total 187 samples from three populations of *Petunia axillaris* subsp. *axillaris* (populations 70 [34° 28’ 00’’ S, 57° 49’ 48’’ W], 71 [34° 30’ 58’’ S, 54° 20’ 44’’ W], and 72 [34° 53’ 17’’ S, 56° 15’ 35’’ W]) and two populations of *P. inflata* (populations 61 [28° 25’ 07’’ S, 58° 57’ 26’’ W] and 62 [29° 00’ 30’’ S, 59° 08’ 26’’ W]) (Ando et al., 1998; Ando et al., 1995). Plants were grown in chambers (16L8D, 22°C) or a greenhouse at the Kashiwanoha campus, Chiba University. Note that our sample set consists of half-sibs of in total 40 families. See Table S4 for details.

### Library preparation and sequencing

We collected styles from flower buds and stored them in RNA later under −80C (De Wit et al., 2012). We extracted total RNA using RNeasy Mini Kit (QIAGEN) and TRI Reagent (Molecular Research Center, Inc) following the manufacturing protocols. We performed the multiplexed sequencing using the PCR-cDNA Barcoding Kit (SQK-PCB109; Oxford Nanopore Technologies). Libraries were prepared according to the manufacturing protocol (PCB_9092_v109_revD_10Oct2019), and 12 barcodes were used per library. We used the MinION device and flow cells R9.4.1 for sequencing. Three libraries were sequenced per flow cell, by washing the flow cell after each run with the Flow Cell Wash Kit (EXP-WSH003 or EXP-WSH004).

### Identification of *S-RNase*

We first trimmed barcode and adaptor sequences using Porechop (https://github.com/rrwick/Porechop; 3 May 2024). We then filtered low-quality reads (Q value <= 6) and trimmed the first 50 bp of 5’ ends using Nanofilt (De Coster et al., 2018). We searched for *S-RNase*-like sequences among those filtered sequences as follows. First, we converted filtered fastq files to fasta files by seqkit (Shen et al., 2016). Then we prepared a set of query sequences that are publicly available 37 *S-RNase* sequences (Table S7). Using these queries, we searched for *S-RNase*-like sequences using blastn (v.2.2.31) (Altschul et al., 1990) and HMMER (v.3.3.2) (Finn et al., 2011) with default parameters. We used sequences that were detected in both blastn and HMMER for the following analyses.

As *S-RNase* alleles should be heterozygous in each individual, we separated them by clustering analysis based on sequence similarity. For each individual, *S-RNase*-like sequences were aligned and distance matrix between sequences was calculated by mafft (Katoh et al., 2002). Using the obtained matrix, we performed hierarchical clustering using the R library dendextend (Galili, 2015). We classified sequence clusters based on the threshold value of 0.7, and omitted clusters to which less than 10 sequences belonged. We realigned sequences within each cluster, and omitted low-frequency indels (< 0.05) as they are likely to be sequence errors by a custom R script. Finally, we obtained a consensus sequence for each cluster using EMBOSS cons (Rice et al., 2000). For each sample, this clustering analysis was performed and *S-RNase*-like sequences were obtained.

We then combined all the obtained *S-RNase*-like sequences, previously reported *S-RNase* sequences of *Petunia*, and those of *Antirrhinum* as outgroups, and generated an alignment using mafft (v.7.310, parameters: --adjustdirectionaccurately). After excluding two *S-RNase*-like sequences that are unlinked to self-incompatibility (*Panon-S* and *PiRNX2*; (Lee et al., 1992)) as well as sequence clusters that have less than 50 reads or 500bp in total length, a maximum likelihood tree was generated using the software IQ-TREE (v.2.1.4-beta, parameters:-m MFP -AIC -B 1000) (Nguyen et al., 2015). Note that we omitted plant samples that had only one *S-RNase*-like sequences other than *Panon-S* and *PiRNX2* satisfying the criteria (number of reads per cluster and sequence length). We classified *S-RNase* based on the monophyly in the tree and assigned S-alleles for each individual. We note that it is not trivial to determine the threshold for delimiting *S*-alleles. Although we confirmed that the tree-based determination was mostly concordant with the pair-wise genetic distance between sequences (Fig. 2b) and that genotypes of S-alleles were completely linked with SI phenotypes as long as investigated in this study (Fig. 5), there might be a few discordances between assigned genotypes and SI specificities.

### Sanger sequencing and PCR-based genotyping

To evaluate the sequencing accuracy of Nanopore-based genotyping, we performed Sanger sequencing for each *S-*allele and compared them with Nanopore-based sequences. We designed primers to amplify *S-RNases* based on the Nanopore sequences (Table S8), and performed PCR using Tks Gflex DNA Polymerase (Takara) according to the manufacturing protocol.

To estimate allele frequency in a larger sample set, we also performed PCR-based genotyping for in total of 187 individuals including those not used for Nanopore sequencing (Table S4). To test the validity of PCR-based genotyping, we performed Sanger sequencing of multiple copies that were assigned as the same *S-*alleles, and generated phylogenetic trees for those *S*-alleles by maximum likelihood method using the IQ-TREE (v.2.1.4-beta, parameters:-m MFP -AIC -B 1000) (Nguyen et al., 2015). While PCR-based genotyping detected two *S-*alleles per diploid heterozygous individual in most cases, we observed false positive amplifications in a few individuals (detailed in Tables S4, S5 and, S8). For those amplifications, we validated PCR fragments by Sanger sequencing, confirming that they are mis-amplification of other *S*-alleles detected in the same individuals (Tables S4, S5 and, S8).

### Estimation of the number of *S*-alleles in each population

The total numbers of *S*-alleles were estimated by using two methods: the *E*_2_ estimator proposed by (O’Donnell and Lawrence, 1984) and a curve fitting using a two-parameter Michaelis-Menten model (Busch et al., 2010). We employed these methods because both of them do not assume equal frequencies of *S*-alleles. For the method of Busch et al. (2010), the estimates were based on the average number of unique *S*-alleles inferred from 1,000 times of resamplings without replacement. For the curve fitting, we used the drc package (Ritz and Streibig, 2005) in R (R Development Core Team 2023). In the Michaelis–Menten model, *f* (*x*) = *S*_max_ / (1 + *K*/*x*), *S*_max_ corresponds to the total number of alleles expected in the population, which was reported for each population.

### Test of association with self-incompatibility phenotypes

To evaluate our method to detect *S*-alleles based on the sequence similarity of the *S-RNase*, we examined whether self-incompatibility phenotypes are associated with *S-RNase* genotypes by crossing experiments. We observed the elongation of pollen tubes in crosses within the same *S*-alleles and between the *S*-alleles. For crossing experiments, we used individuals of *P. axillaris* subsp. *axillaris*, while both *P. axillaris* subsp. *axillaris* and *P. inflata* were used for sequencing analysis.

We performed two sets of crossing experiments. For the first one, seven homozygous lines generated by forced selfing were used as pollen donors (*S*7, *S*22, *S*17, *S*3, *S*g, *S*f, and *S*j), and heterozygous individuals that have at least one of these *S*-alleles were used as pistil donors (Table S6). For the second ones, we used homozygous lines of *S*6, *S*17, and *S*d alleles as pollen and pistil donors. These three *S*-alleles (*S*6, *S*17, and *S*d) had closely related *S-RNase* sequences.

We observed pollen tubes by aniline staining as follows. We first removed anthers from flower buds and closed them by staplers to avoid contamination. Styles were pollinated 24 hours after emasculation and harvested 48 hours after pollination. Harvested styles were fixed in a 3:1 mixture of ethanol and acetic acid, softened for 8 hours in 1M NaOH at 65 °C and stained with aniline blue in a 2% K_3_PO_4_ solution overnight at room temperature. Styles were mounted on slides to examine the pollen tubes using epifluorescence microscopy (Olympus BX53). We counted the number of pollen tubes that reached the bottom tip of styles. Less than 20 pollen tubes were considered as a criterion of incompatible crosses.

### Phylogenetic analysis with *S-RNase* of other genera

To understand the phylogenetic relationship between *Petunia S-RNase* obtained in this study and previously reported *S-RNase* including those from other genera of Solanaceae, we constructed a phylogenetic tree by generating a large sequence alignment. The list of sequences used for the tree is available in Table S7. A phylogenetic tree was generated by the maximum likelihood method using the IQ-TREE (Nguyen et al., 2015).

### Detection of positively selected sites and sliding window analysis

To detect amino acid positions under selection in the *S-RNase* gene from *Petunia*, we estimated non-synonymous/synonymous substitution ratio for each site by using the PAML4 software (Yang, 2007). The codeml package of PAML4 was used to calculate posterior probabilities of codon sites under positive selection with the Bayes empirical Bayes (BEB) method. The likelihood of two models M1a (nearly neutral) and M2a (positive selection) were compared by a likelihood ratio test. For this analysis, we used in total 79 sequences of *Petunia* obtained by Sanger sequencing or by previous studies (Table S7). A phylogenetic tree used as an input for PAML4 was generated by a maximum likelihood method with the General Time Reversible model using MEGA11 (Tamura et al., 2021).

To evaluate the genetic diversity along the *S-RNase* gene from *Petunia*, we performed sliding window analysis using DnaSP v. 6.12.03 (Rozas et al., 2017). The window size was 50 and the step size was 10. Used sequences were mostly the same as those used for PAML4, but we omitted a few relatively short sequences due to designed primer positions (Tables S7 and S8).

## Acknowledgments

This work was supported by the FOREST program of the Japan Science and Technology Agency (JPMJFR2046 to TT), Grants-in-Aid for Scientific Research (18H04813, 19H03271, 20H04856, 22K21352, 23H02537 to TT, 21K06079 to KK and 17H04605 to KU), Toray Science and Technology Grant and Inamori Research Grant to TT.

## Data availability

Sequence data have been deposited in DDBJ under accession numbers LC819163-LC819242.

