## Supplementary Figure 1 for "Long-read sequencing reveals the allelic diversity of the self-incompatibility gene across natural populations in *Petunia* (Solanaceae)"

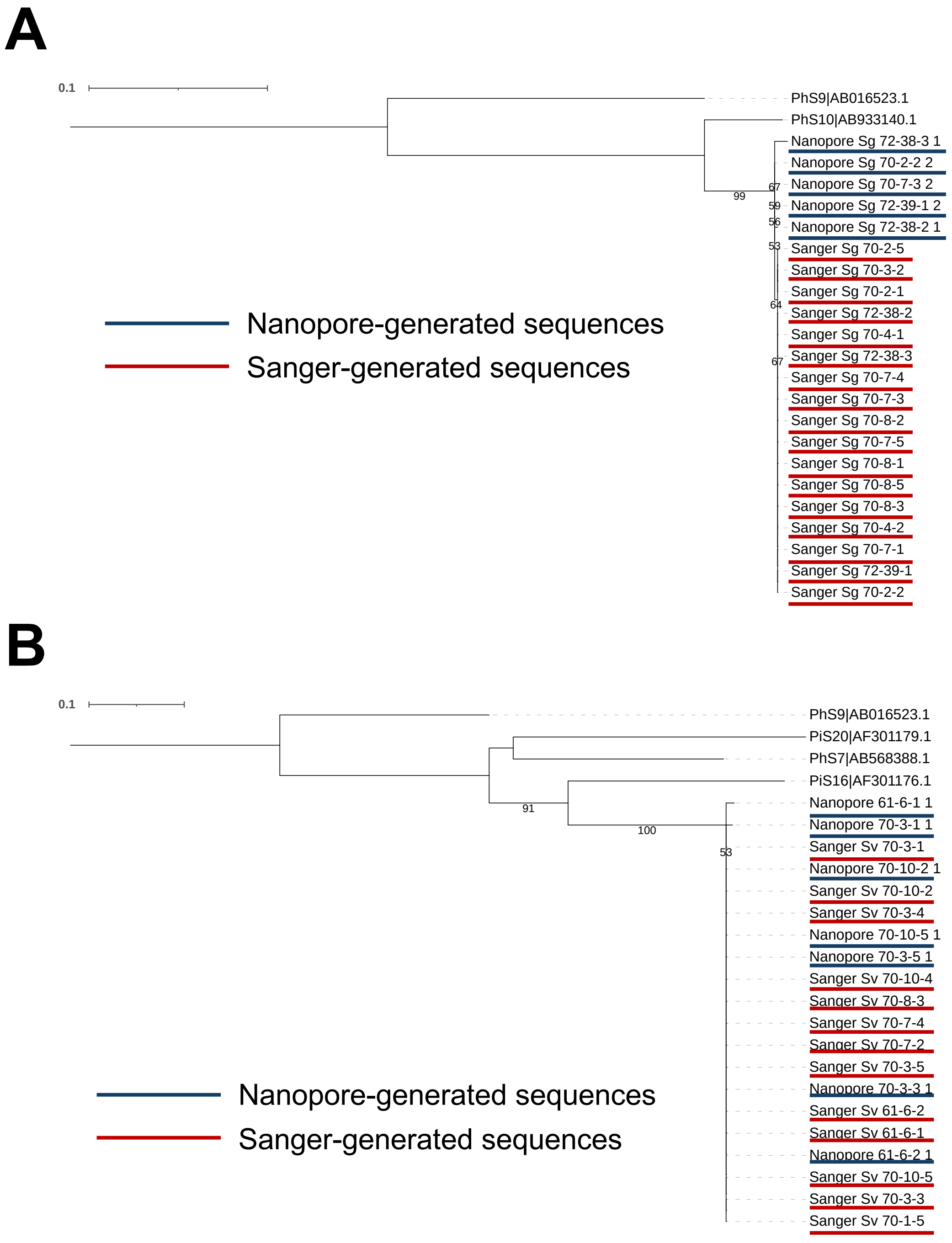


**Fig. S1**. Maximum likelihood phylogenetic trees of *S-RNase* within the *S*-alleles, *S*g (A) and *S*v (B). Nanopore and Sanger-generated sequences are indicated by blue and red lines, respectively. The values on the branches indicate the percentage of 1,000 bootstrap replicates.
